# Intraguild predation does not necessarily have negative effects on pest biological control: insights from a multiple consumer–multiple resource food web population model

**DOI:** 10.1101/2021.08.09.455696

**Authors:** Michel Iskin da S. Costa, Lucas dos Anjos

## Abstract

We develop a food web population dynamical based on an experimental pest biocontrol setup consisting of thrips and aphids (pests) being consumed by two agents *Macrolophus pygmaeus* and *Orius laevigatus*, and with *O. laevigatus* being an intraguild predator of *M. pygmaeus*. By means of numerical simulations, we show that pest biocontrol disruption can be avoided depending on initial population densities of pests and agents, despite the intraguild predation (IGP) of *O. laevigatus* upon *M. pygmaeus*. This possible avoidance of pest biocontrol disruption is in accordance with the referred experimental setup and moreover, the proposed model corroborates the importance of initial densities of pests and agents in the determination of the failure or success of pest biocontrol found in this and other biocontrol experimental setups.

## Introduction

Generalist predators that feed on more than one species of prey have proven to be efficient biological control agents (e.g., Gardiner and Landis, 2007). Because most crops are attacked by more than one species of pest, biological control programs, especially in greenhouse crops, are increasingly based on releases of generalist predators against common greenhouse pests such as thrips, whiteflies, spider mites, aphids, and leaf miner moths (Messelink, 2012).

Despite their broad diet spectrum, generalist predators do not always control all pests (Symondson et al., 2002) and other natural enemies are needed in such situations. One approach is to release several species of generalist predators for multiple pest control. However, generalist predators are often involved in the competition for shared prey and predation upon each other (intraguild predation, IGP (Polis et al. (1989))), which can affect both their coexistence and the results of biological control.

In order to assess whether the negative effects of IGP on pest biocontrol can be mitigated in a specific biological setup, Messelink and Janssen. (2014) evaluated in an experimental study the coexistence of two generalist predatory bugs *Macrolophus pygmaeus* Rambur (Hemiptera:Miridae) and *Orius laevigatus* (Fieber) (Hemiptera: Anthocoridae) in a sweet pepper crop with two pest species as shared resources. The two pests used were the peach aphid *Myzus persicae* (Sulzer) (Hemiptera: Aphididae) and the western flower thrips *Frankliniella occidentalis* Pergande (Thysanoptera: Thripidae), both important pests in sweet pepper. Moreover, they also found a unidirectional intraguild predation for *Orius majusculus* (Reuter) (Hemiptera: Anthocoridae) preying on *M. pygmaeus*.

Their study shows that despite being involved in intraguild predation, the two agents (predators) complemented each other in the control of the two pests.

Motivated by the experimental finding that these two agents (predators) can coexist in a sweet pepper crop, and their presence does not affect the control of the two pest species we developed a population dynamical model that can qualitatively generate this lack of pest control disruption found in the laboratory experiments carried out by Messelink and Janssen (2014).

The outline of the present work is as follows. In section 2 we present the model together with the interpretation of its variables and parameters. In section 3 we perform the numerical bifurcation analysis of the model and present the biological results in terms of pest biocontrol. In section 4 we discuss the results of the present work.

## 2 Methods

### A multiple consumer–multiple resource food web model

Messelink and Janssen (2014) performed experiments to evaluate the co–occurrence of the generalist predators *Macrolophus pygmaeus* and *Orius laevigatus* and their control of two pests in a sweet pepper crop. Both predators prey on thrips and aphids, and *O. laevigatus* is an intraguild predator of *M. pygmaeus*. Their experimental food web is schematically shown in figure 1.

**Figure 1.**
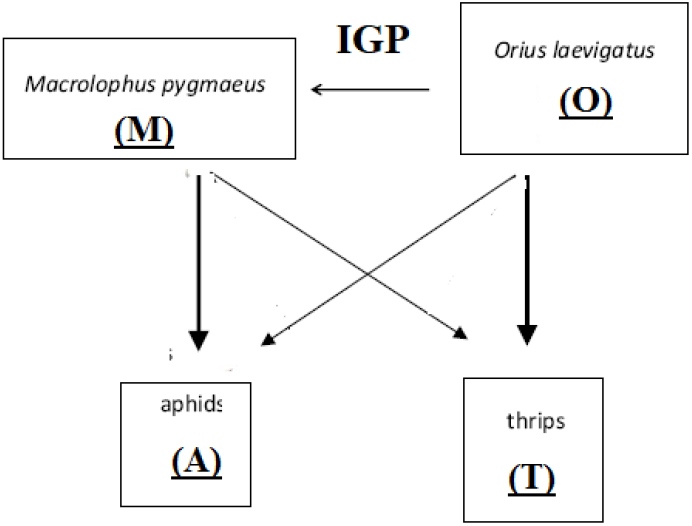
A food web containing multiple consumer–multiple resource interactions according to the experiment performed in Messelink and Janssen (2014). Arrows indicate predation. IGP means intraguild predation.

A continuous–time mathematical population dynamical model of the species in the food web depicted in figure 1 can take on the form:

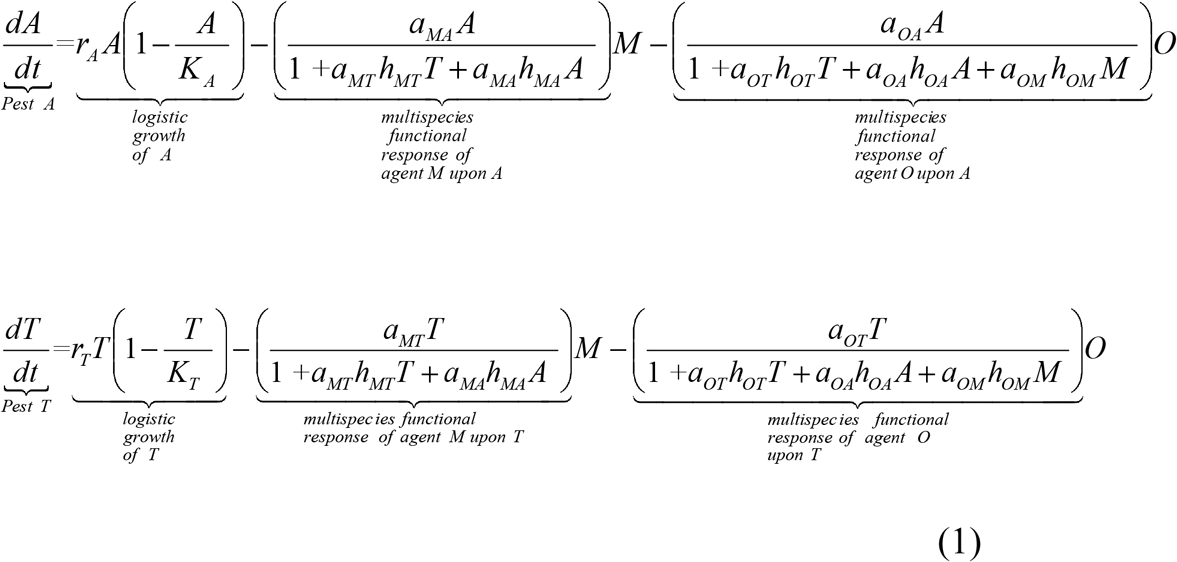

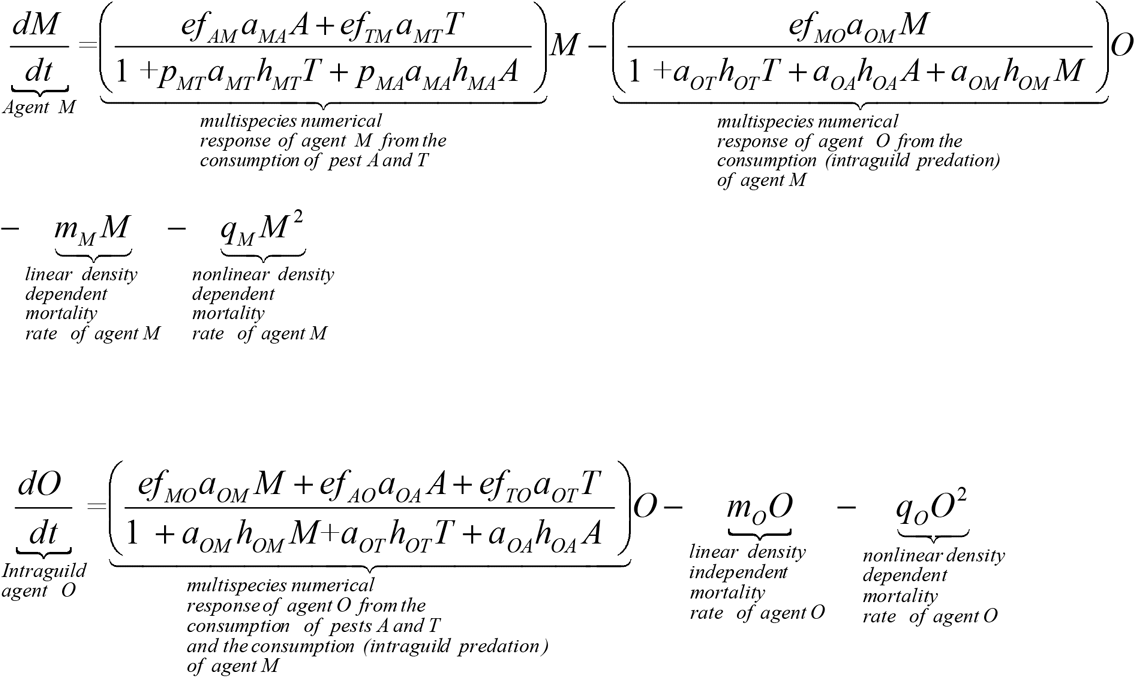

*A* and *T* are the densities of the pests (prey), while *O* and *M* represent the densities of the control agents (predators). The definitions of the variables and the parameters of the model (1) are shown in table 1.

**Table 1.**
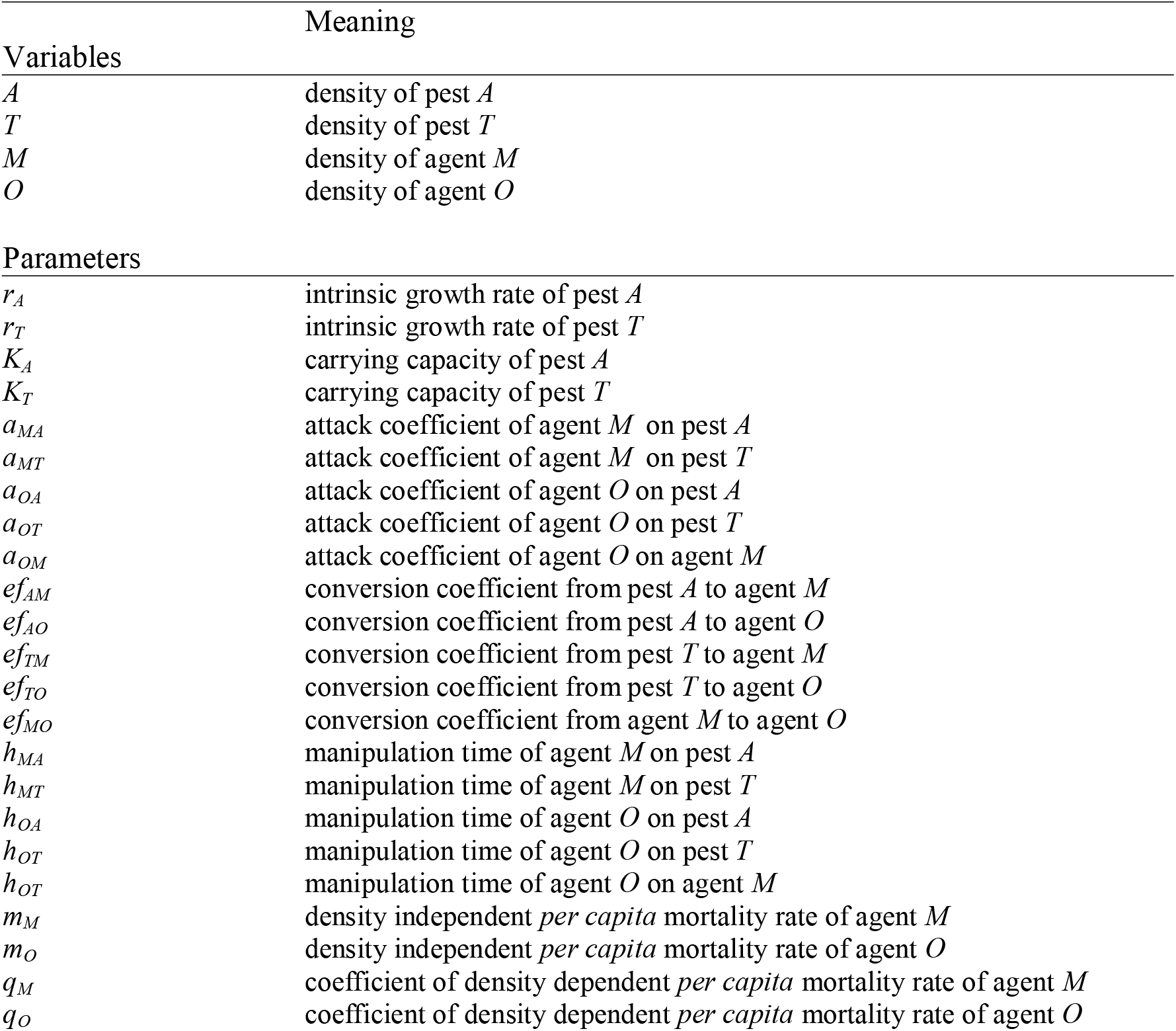
Definition of variables and parameters of the model (1)

We assume a logistic growth in the pests *A* and *T* because of the presumed lack of exploitative competition among them. This assumption is based on another experiment where exploitative competition between the pests was improbable, certainly due to the large leaf size of the crop (Messelink et al., 2008).

The density dependent–mortality rate of the agents *M* and *O*, i.e., the expressions *q_M_M*^2^ and *q_O_O*^2^ can be associated with con–specific cannibalism. For instance, control agent such as spider mite of the species Amblyseius *swirski AthiasHenriot* (Acari:Phytoseiidae) is subject to con–specific cannibalism in the early stages of their life cycle (Rasmy et al., 2004).

In Messelink and Janssen (2014) it is experimentally shown that besides the coexistence of the two agents, the intraguild predation between these agents does not disrupt pest biological control. Therefore, regarding this experimental result, it is interesting to investigate if our proposed model (1) can yield this same result in qualitative terms. This can be verified by increasing the value of the intraguild predation attack coefficient of *O. laevigatus* upon *M. pygmaeus* (*a_OM_*) in the model (1) and checking how this increase in IGP affects the densities of all populations in the equilibrium in the model (1). By having the values of the population densities we can assess whether the agents coexist and do not disrupt biocontrol (as found in the experiment). To perform this analysis, a numerical bifurcation of the model (1) as a function of *a_OM_* by means of the software package XPPAUT (Ermentrout, 2002) is undertaken. In essence, this software calculates the equilibrium points of the nonlinear differential equations given by the model (1) (i.e., the numerical solutions to the system of equations given by *dA*/*dt*=0, *dT*/*dt*=0, *dM*/*dt*=0, and *dO*/*dt*=0) as one varies, for instance, the parameter *a_OM_*, together with the real part of their corresponding eigenvalues. This information is gathered to draw the graphs with equilibrium population density levels and their respective stability characteristics (stable/unstable equilibrium points) displayed throughout this work. Moreover, we call attention to the fact that the choice of the parameter values of the model (1) was partially guided by an intention to create, when possible, stable dynamics in the analyzed models, avoiding thus, for instance, sustained oscillations (e.g., limit cycles). In this way, all species densities variation of the model (1) as a consequence of changes in the intensity of the intraguild coefficient attack *a_OM_* can be promptly read off the bifurcation diagrams without resorting to mean value calculation of the species densities due to the species temporal oscillations.

## Results

The resulting bifurcation diagram of all the involved species (*A, T, M, O*) of the model (1) is displayed in figure 2. These diagrams show the densities of the species when the intraguild predation attack coefficient of *O. laevigatus* upon *M. pygmeus* (*a_OM_*) is increased.

**Figure 2.**
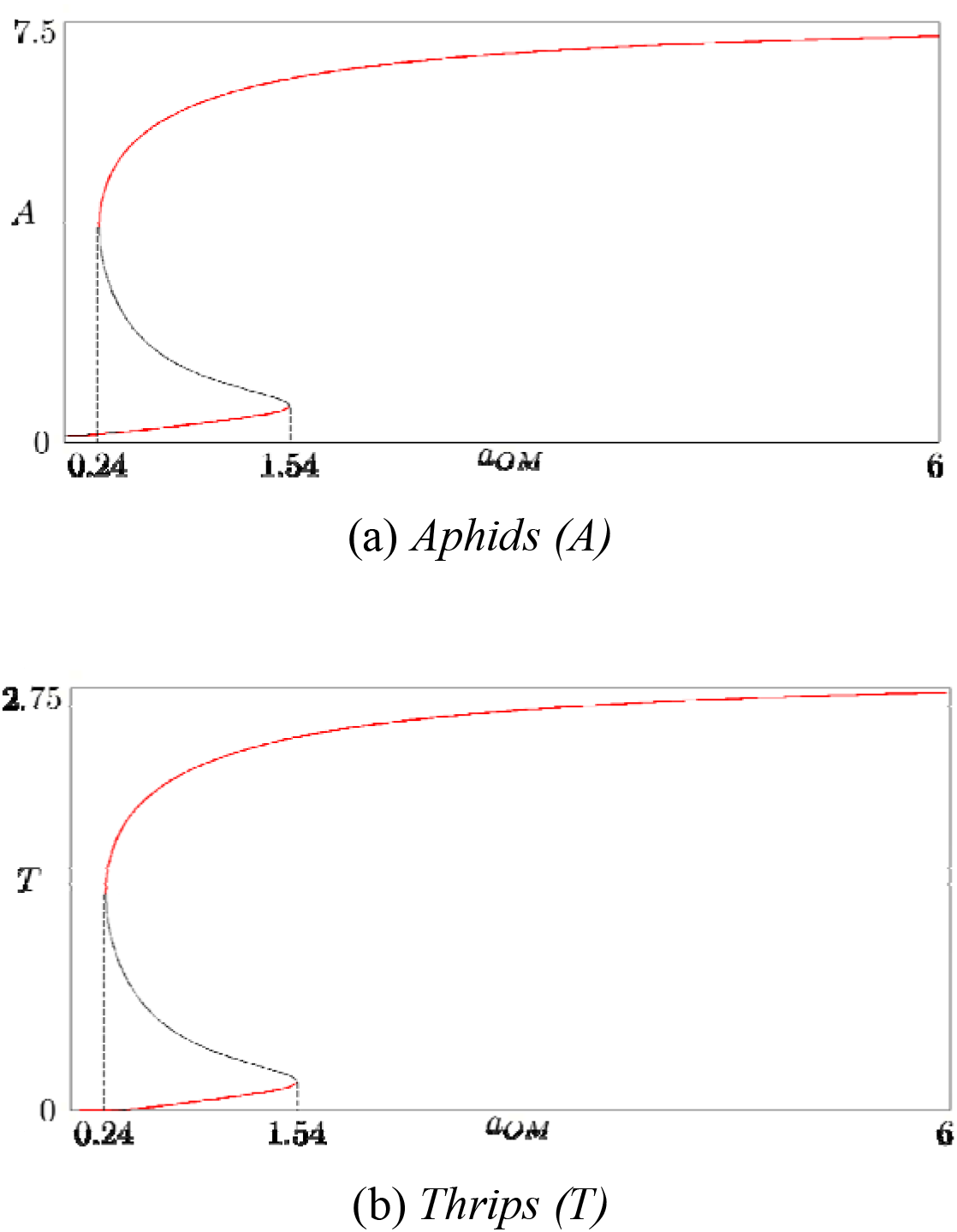
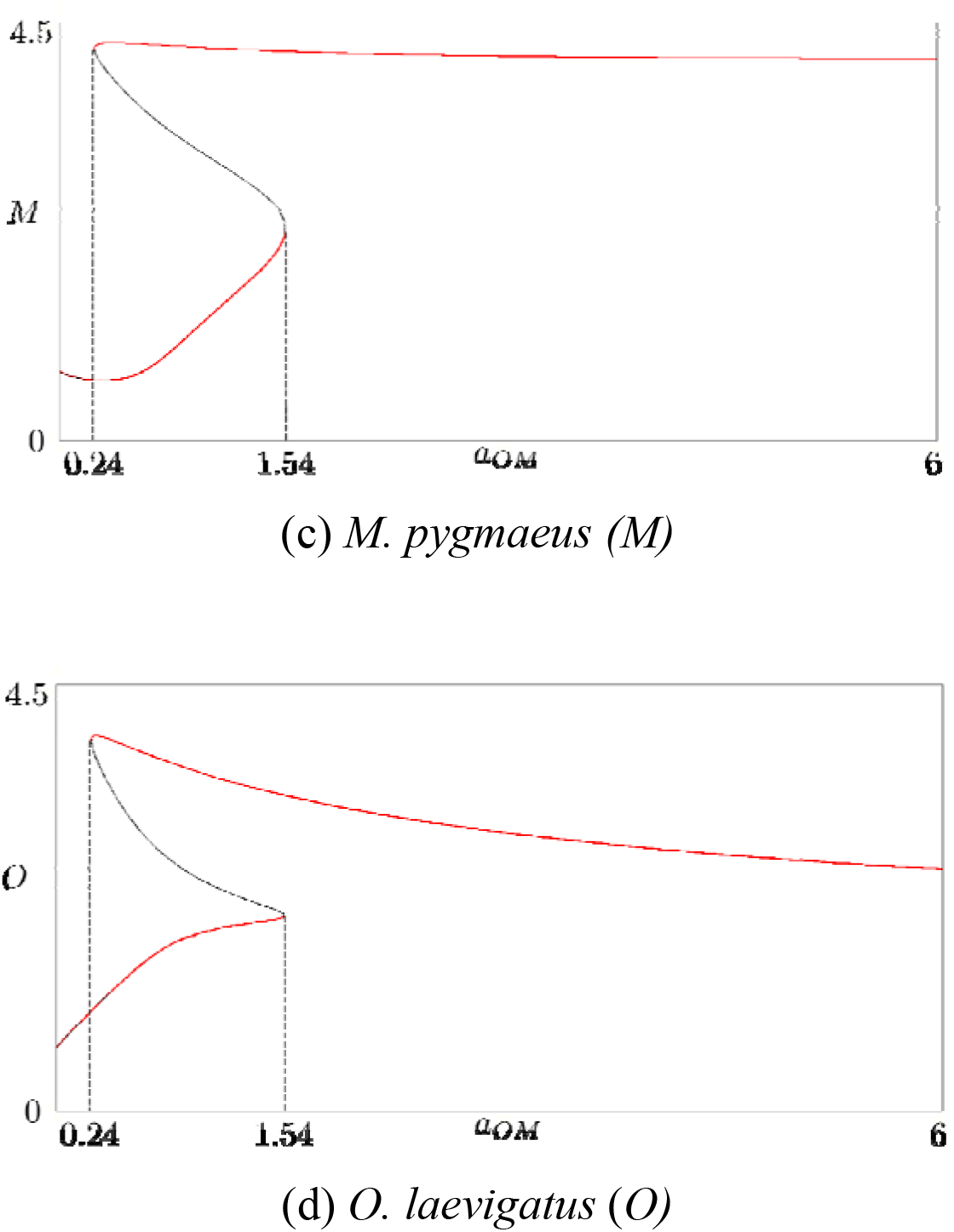
A numerical bifurcation diagram of model (1) as a function of the intraguild predation attack coefficient of *O. laevigatus* upon *M. pygmaeus* (*a_OM_*). Bistabiliy occurs along the interval 0.24 < *a_OM_* < 1.54. Red lines: stable equilibrium points; black lines: unstable equilibrium points. Parameter values: *r_A_*=1.12; *r_T_*=1.1; *K_A_*=9.12; *K_T_*=3.444; *a_OT_*=0.9; *a_OA_*=0.9; *a_MT_*=1; *a_MA_*=1; *ef_MO_*=5; *ef_AO_*=5; *ef_TO_*=5; *ef_TM_*=2; *ef_AM_*=2; *h_OM_*=2; *h_OT_*=2; *h_OA_*=2; *h_MT_*=2; *h_MA_*=2; *m_T_*=0.1; *m_O_*=0.1; *q_M_*=0.5; *q_O_*=0.5.

Note first that the food web in figure 1 consists of two community modules (Holt, 1997) of type “two prey species that share one predator” (i.e., *A* – *T* – *M* and *A* – *T* – *O*) which are connected also by the intraguild predation of *O* upon *M*(i.e., coefficient *a_OM_*). These two community modules (*A* – *T* – *M* and *A* – *T* – *O*) are known to possess positive bistability (Abrams and Matsuda, 1996) (here, positive bistability is the occurrence of two different population levels in which all species can coexist). Actually, the choice of the parameter values was also guided by the creation of a positive bistability in the model (1) (as shown in figure 2) on account of the following reason: this bistability in the model (1), which occurs throughout the interval 0.24 < *a_OM_* < 1.54, can provide a two-fold explanation: (i) the red lower branch of the pests *A* and *T* may well represent pest suppression (relatively low pest levels), while (ii) the red upper branch of the pests *A* and *T* may well represent the disruption of pest control (relatively high pest levels). In a way, item (i) corroborates that predation of one agent upon the other (the intraguild predation of *O. laevigatus* upon *M. pygmaeus*) does not necessarily have negative effects on biological control of aphids and thrips (Messelink and Janssen, 2014).

Importantly, model (1) also suggests that these two outcomes in pest control (pest suppression and disruption of pest control) depend on the initial level of the populations (a consequence of the positive bistability of the model (1); see figure 2). This result corroborates the importance of initial population densities in pest control (Messelink and Janssen (2014), p.4; see also observations about the importance of initial population densities in a biocontrol prey–predator experimental setup in Leman and Messelink (2014)). However, to be more precise with respect to the mentioned texts in the above references, one should carry out time–series simulations of the model (1) with varying initial densities of the predators *O. laevigatus* (i.e., *O*(0)) and *M. pygmaeus* (i.e., *M*(0)) and check the final densities of the pests *Aphids* (*A*) and *Thrips* (*T*) so as to assess the efficiency of pest control with respect to initial densities of agents (e.g., inoculative (low levels of) /inundative (high levels of) agent releases). Nonetheless, it is important to remark that the model (1) also suggests that increasing the intraguild predation attack coefficient of *O. laevigatus* upon *M. pygmaeus* (*a_OM_*) brings about the increase of both pest levels in the lower and in the upper branch of the bifurcation diagrams of figure 2.

## Discussion

Messelink and Janssen (2014) experimentally evaluated how the co–occurrence of the generalist control agents *Macrolophus pygmaeus* and *Orius laevigatus* affected their control of two pests in a sweet pepper crop. Both agents prey on thrips and aphids, and *O. laevigatus* is an intraguild predator of *M. pygmaeus*. Their study provides further evidence that the use of natural enemies that can be involved in intraguild predation does not necessarily have negative effects on biological control (confirming other experimental studies). Moreover, they also mention that the mechanisms that prevent such negative effects of intraguild predation and the exclusion of one agent (predator) by the other remain elusive.

In this work, by means of simulations of a proposed theoretical food web population dynamical model describing the biotic species interactions in the mentioned experiment, we showed that the use of natural enemies that can be involved in intraguild predation does not necessarily have negative effects on biological control. This result is conveyed in the model (1) by the relatively low pests’ equilibrium densities in figure 2. Furthermore, from a myriad of possible biological processes, we narrowed them down to those included in the model (1) as potential candidates to explain to some extent how negative effects of intraguild predation and the exclusion of one agent (predator) by the other can be avoided in this specific intraguild food web. That is to say, perhaps conceptual (strategic/phenomenological) models (May, 2001) such as model (1) (and/or others) could shed some light to help unveil these mechanisms – at least, qualitatively– and thereby lessen their seeming elusiveness in experimental setups.

In a more general context of food web population dynamics theory, we investigated how the effects of a disturbance (in our case, an increase in the intensity of the intraguild attack coefficient *a_OM_*) propagate through the species densities of a specific food web. We think that this conjunction of applied and theoretical ecology can contribute to expanding the understanding of how natural enemy–density–mediated indirect interactions may be used to enhance pest biocontrol strategies (Chailleux et al. 2014).

